# Human ES and iPS Cells Display Less Drug Resistance Than Differentiated Cells, and Naïve-State Induction Further Decreases Drug Resistance

**DOI:** 10.1101/2020.08.17.253492

**Authors:** Yulia Panina, Junko Yamane, Kenta Kobayashi, Hideko Sone, Wataru Fujibuchi

## Abstract

Pluripotent stem cells (PSCs) possess unique characteristics that distinguish them from other cell types. Human embryonic stem (ES) cells are recently gaining attention as a powerful tool for human toxicity assessment without the use of experimental animals, and an embryonic stem cell test (EST) was introduced for this purpose. However, human PSCs have not been thoroughly investigated in terms of drug resistance or compared with other cell types or cell states, such as naïve state, to date. Aiming to close this gap in research knowledge, we assessed and compared several human PSC lines for their resistance to drug exposure. Firstly, we report that RIKEN-2A human induced pluripotent stem (iPS) cells possessed approximately the same sensitivity to selected drugs as KhES-3 human ES cells. Secondly, both ES and iPS cells were several times less resistant to drug exposure than other non-pluripotent cell types. Finally, we showed that iPS cells subjected to naïve-state induction procedures exhibited a sharp increase in drug sensitivity. Upon passage of these naïve-like cells in non-naïve PSC culture medium, their sensitivity to drug exposure decreased. We thus revealed differences in sensitivity to drug exposure among different types or states of PSCs and, importantly, indicated that naïve-state induction could increase this sensitivity.

## Introduction

Pluripotent stem cells (PSCs) possess unique characteristics that distinguish them from other cell types. They are able to self-renew, differentiate into all three germ layers, and produce a wide variety of differentiated cell types (Keller, 2005; Martin, 1981). Embryonic stem (ES) cells are recently gaining attention as a tool to investigate toxicity without the use of living animal experiments, and the *in vitro* (mouse) embryonic stem cell test (EST) was developed specifically for this purpose (Liu et al., 2017a; Tandon and Jyoti, 2012). The human EST (hEST) shortly followed, and its successful application to developmental toxicity prediction was reported (Adler et al., 2008; Yamane et al., 2016). However, the use of human ES cells has raised important ethical issues, and the replacement of hEST with an analogous induced pluripotent stem (iPS) cell test is highly desired. To develop such tests, it is necessary to study iPS cells and their drug sensitivity in comparison with ES cells.

Stem cells have been investigated extensively for drug sensitivity and drug resistance, and compared with non-stem cells in cancer research (Gottesman et al., 2002). Cancer stem cell studies suggest that stemness confers resistance to drug exposure in many, if not all, cancers, including melanoma (Fukunaga-Kalabis and Herlyn, 2012; Luo et al., 2012; Wouters et al., 2013), ovarian cancer (Hu et al., 2010), pancreatic cancer (Li et al., 2015; Niess et al., 2015), lung cancer (MacDonagh et al., 2016; Yeh et al., 2012), colorectal cancer (Tsunekuni et al., 2019; Zhou et al., 2018), and many others. In cancer, this resistance is called multidrug resistance and is a widely known feature of stem cells (Gillet and Gottesman, 2010; Gottesman et al., 2002). Multidrug resistance in cancer stem cells is thought to result from the increased expression of drug-efflux genes (Ambudkar et al., 1999), reduced drug uptake (Shen et al., 2000), blocked apoptosis (Gottesman et al., 2002), and possibly other mechanisms, which are thought to be the features of cancer stem cells. On the other hand, scarce information on the drug sensitivity of non-cancer stem cells suggests that stem cells are more sensitive to drug exposure than differentiated cells (Kusakawa et al., 2008; Laschinski et al., 1991). To resolve this controversy, more studies on the drug sensitivity of non-cancer stem cells are needed. However, work comparing the drug sensitivity of different human PSCs is lacking.

Recent works in the stem cell field have uncovered the existence of a variety of pluripotent states. First, in 2007, it was discovered that human ES cells *in vitro* differed from mouse ES cells derived from preimplantation blastocyst (Brons et al., 2007; Tesar et al., 2007). Human ES cells resemble the more differentiated mouse epiblast stem cells, also called “primed ES cells”, rather than the less differentiated blastocyst-derived “naïve ES cells”. Thus, the notion of different pluripotent states was put forward and researchers began to seek ways of naïve-state induction in human ES cells. The subsequent rapid appearance of different methods of naïve-state induction (Carter et al., 2016; Chan et al., 2013; Chen et al., 2015; Duggal et al., 2015; Gu et al., 2012; Hanna et al., 2010; Li et al., 2009; Panova et al., 2018; Park et al., 2018; Qin et al., 2016; Takashima et al., 2014; Valamehr et al., 2014; Wang et al., 2011; Ware et al., 2014; Warrier et al., 2016; Zimmerlin et al., 2016) and advances in RNA sequencing techniques revealed that different methods produce different “states” of pluripotency, with several “intermediate” states existing on the naïve-primed axis (Liu et al., 2017b; Warrier et al., 2017). Human iPS cells have for a long time existed in the gray area, and their equivalent status to ES cells has been hotly debated (Christodoulou and Kotton, 2012; Hackett and Fortier, 2011; Narsinh et al., 2011). In particular, iPS cell cultures appear more heterogeneous than ES cell cultures (Hotta et al., 2009a; Takahashi and Yamanaka, 2006) and may require a selection of “best-quality” cells, suggesting that iPS cell lines may contain cells with different stemness properties. As the link between stemness properties and drug resistance properties is important for both basic research and toxicology research, more work on the direct comparison of the general drug resistance of human ES cells and iPS cells is needed.

In this work, we investigated the sensitivity of different human PSCs to drug exposure. Our goals were to 1) compare the drug resistance properties of ES cells and non-pluripotent cell types, 2) compare the drug resistance of human ES cells and human iPS cells, and 3) determine whether naïve-state induction methods, such as EOS selection and application of existing naïve-state induction medium, would increase or decrease the sensitivity of pluripotent cells to drug exposure.

## Materials and methods

### Drugs

The following drugs were used for toxicity assessment in this study: amiodarone (TCI, Tokyo, Japan, A2530-1G), atorvastatin (TCI, Tokyo, Japan, A2476-1G), clotrimazole (Wako, Tokyo, Japan, 035-16021), aspirin (Wako, Tokyo, Japan, 015-10262), cyclosporin A (Wako, Tokyo, Japan, 031-24931), chlorpheniramine (Wako, Tokyo, Japan, 030-13271), chlorpromazine (Wako, Tokyo, Japan, 033-10581), and ibuprofen (Wako, Tokyo, Japan, 098-02641). All drugs were diluted with DMSO to the desired concentrations.

### Cell culture

ES cells and iPS cells were routinely cultured in StemFit AK02N (Ajinomoto, Tokyo, Japan), which is one of the standard media for iPS cell culture, e.g. (Kubara et al., 2018), with the addition of 10 μM ROCK inhibitor Y-27632 (CultureSure, Wako, Tokyo, Japan, 030-24023) and 1 μL/mL iMatrix-511 (Nippi, Tokyo, Japan, 892014). Routine splitting was performed by washing with PBS (room temperature), addition and subsequent removal of TrypLE Express Enzyme (ThermoFisher, Tokyo, Japan, 12604013), gathering the cells with the AK02N medium through the addition of 10 μM Y-27632, centrifuging at 500 rpm for 5 min, aspiration of the medium, resuspending the cells in new medium, and plating at the required density. All routine cultures were performed in 6-well plates (Greiner, 657160, Merck, Darmstadt, Germany) at 37°C in 5% CO_2_, and splitting was performed at 80% confluence.

### ATP assay

The ATP assay was used to assess and compare the sensitivity of the cell lines to drug exposure. We define cell sensitivity as the decrease in viability of the cells relative to control sample (vehicle-treated cells under the same conditions). For the ATP assay, cells were seeded at the final concentration of 8,000 cells/well onto Falcon® 96-well black clear bottom flat-bottomed plates (353219) in the total volume of 100 μL of medium per well. The cells were then incubated for 24 hours at 37°C in 5% CO_2_. The next day, the medium was changed to a new medium containing different doses of the drugs under investigation. All drugs used in this study were diluted with DMSO (vehicle), and the final concentration of DMSO in each well was always 0.1% (including control sample). After exposure to the drugs, the cells were cultured for 24 hours at 37°C in 5% CO_2_, and CellTiter-Glo® Luminescent Cell Viability Assay (Promega, Tokyo, Japan, G7571) was used for the ATP reading according to the manufacturer’s instructions. The samples were subjected to plate-reading using the Perkin Elmer EnVision 2104 plate reader, PerkinElmer, USA. The reproducibility of the ATP assay was ensured by commissioning two independent operators to conduct the experiment, as separate biological replicates.

### ED10 determination

The extension package “*drc*” (dose-response curve) (Ritz et al., 2015) for the statistical environment R was used to analyze data, determine ED10 values, and plot the dose-response curves for each drug. The *drc* package provides the code for non-linear model fitting. The specification of this dose-response (regression) model describes the mean by a parametric function of dose and the assumptions of the response distribution. In this work, the log-logistic function was used to plot the dose-response curves, and the confidence interval (95%) was calculated and included for each curve. ED10 values were calculated on the basis of curve fitting using the *drc* package.

### EOS introduction and EOS-based cell selection

To select the best-quality iPS cells, the EOS selection system (Hotta et al., 2009a, 2009b) was used. This system takes advantage of the fact that naïve PSCs express Oct3/4 from the distal enhancer (as opposed to the proximal enhancer in primed cells (Choi et al., 2016; Tesar et al., 2007; Yeom et al., 1996)), enabling the selection of high-quality naïve-like iPS or ES cells by GFP expression or antibiotic selection. (PB) EOS-C(3+)-GFP/puroR vector was a kind gift from Professor Akitsu Hotta. To introduce the vector, the cells were cultured to 80% confluence and transfected using QIAGEN Effectene Transfection Reagent (Cat. no. 301425, Tokyo, Japan) as indicated by the manufacturer. The next day after transfection, 0.5 μg/mL puromycin (Sigma, P9620-10ML, Merck, Darmstadt, Germany) was added to the cells. After 5 hours, the medium was changed to 1 μg/mL puromycin medium and dead cells were aspirated. The cells were left overnight and the next day, the medium was changed again and dead cells were aspirated. As a result, 2–3 colonies consisting of 5–7 cells remained, and these were subsequently expanded, their puromycin resistance confirmed by applying 5 μg/mL puromycin to the culture medium.

### Naïve-state induction methods

First, we surveyed the literature for the most recent and well-established methods for naïve-state induction. As a result, we chose four methods, namely, 5i/LAF method (Theunissen et al., 2014), t2iLGoY method (Takashima et al., 2014), YAP method (Qin et al., 2016), and XAV939 method (Zimmerlin et al., 2016). We assembled the required media (as shown in pink in Sup. Fig. 1) and added the required additives (such as Y27632) in the quantities recommended in the original articles at designated times during the culture. The cells were cultured in 6-well plates at 37°C in 5% CO_2_, as per instructions in the respective publications. However, we did not use feeder cells (contrary to the instructions in the publications, all of which performed naïve-state induction on feeder cells). The cells were assessed for shape change and stained for the presence of alkaline phosphatase (AP, marker of pluripotency) after naïve-state induction should be complete as indicated in the respective publications.

### Modified YAP method

For the development of the modified YAP method, we first surveyed the literature to find out the effect of each of the additive ingredients. The t2iLGoY method was excluded because it required transfection and overexpression of Nanog gene. The 5i/LAF method and the XAV939 method were excluded because they required the presence of fibroblast growth factor (FGF), which is known to induce differentiation in naïve mouse ES cells, leading them to transition from the naïve state to the primed state (Greber et al., 2010; Ochiai et al., 2015). Thus, we chose the YAP method for the development of our original culture method. We subsequently excluded LIF from the recipe because it affected the JAK/STAT signaling pathway that is important for the mouse ES cells when they are in diapause (Onishi and Zandstra, 2015), and because human embryos lack the diapause stage. CHIR99021 was excluded because it drove the beta-catenin signaling pathway that led to the fibroblast phenotype in our cell lines (data not shown). Forskolin was excluded to avoid fluctuations in intracellular cyclic AMP. As a result, an original recipe was developed, which was based on YAP activator lysophosphatidic acid (LPA, Cayman Chemical, Michigan, USA, CAY-62215-1MG) and included standard ROCK inhibitor Y27632 and MEK inhibitor PD0325901 (Wako, Tokyo, Japan, 162-25291). This medium was applied to stable EOS cell lines and yielded round colonies that were transferred and cultured in AK02N medium for further passaging.

### Immunostaining

The immunofluorescence analysis was performed in the following way. Round colonies of EOS-selected, YAP-treated cells were placed on 30-mm glass bottom dishes (MATSUNAMI, Tokyo, Japan, D11530H) and grown for 24 hours in YAP medium. On the day of immunostaining, the colonies were washed with PBS, fixed with 4% PFA (Wako, Tokyo, Japan,163-20145) for 15 min at room temperature, and permeabilized with 0.5% Triton (Sigma-Aldrich, X100-5ML, Merck, Darmstadt, Germany) in PBS with 10% FBS addition for 30□min. The following antibodies were applied: primary anti-SUSD2 (Sigma, HPA004117-100UL, Merck, Darmstadt, Germany) and anti-KLF17 (Sigma, HPA024629-100UL, Merck, Darmstadt, Germany), in PBS with 10% FBS addition, for 1□hour at room temperature. After 1 hour, the cells were washed with PBS twice and incubated with secondary antibodies: goat anti-rabbit IgG H&L (Alexa Fluor® 594) (Abcam, Tokyo, Japan, ab150080, 1/500 dilution) in PBS with 10% FBS addition, for 1□hour at room temperature. Then, the cells were washed 4 times with PBS, and 2□mL of PBS per dish was added for imaging. Imaging was performed using a KEYENCE BZ-9000 BIOREVO fluorescent microscope.

### Alkaline phosphatase staining and imaging

For AP staining, the cells were briefly washed with PBS, fixed for 5□min with 4% PFA at room temperature, and stained with Alkaline Phosphatase Kit II (Stemgent, #00-0055, Virginia, USA) according to the manufacturer’s protocol. Imaging was carried out using an Olympus CKX41 inverted microscope.

## Results

### Selection of drugs and their initial analysis

As the first step of our analysis, we chose eight drugs with known hepatotoxicities from the Drug-Induced Liver Injury Rank (DILIrank) public database (Chen et al., 2016). This database is available on the US Food and Drug Administration website (https://www.fda.gov/science-research/liver-toxicity-knowledge-base-ltkb/drug-induced-liver-injury-rank-dilirank-dataset) and contains 1,036 FDA-approved drugs. The drugs in the database are ranked by their potential to cause drug-induced liver injury (DILI) on the basis of the analysis of hepatotoxicity descriptions in FDA-approved drug labeling documents and available medical literature. All drugs in the database are divided into four classes: No-DILI-concern (hereinafter referred to as class I), vLess-DILI-concern (hereinafter referred to as class II), vMost-DILI-concern (hereinafter referred to as class III), and Ambiguous-DILI-concern (in which the exact cause of liver injury could not be ascertained). In addition to vDILI-concern class, each drug is assigned a DILI severity score ranging from 0 to 8. We chose the eight drugs so that those would represent the three classes of vDILI-concern and various DILI severity scores (Table 1). From class I, we chose chlorpheniramine, which has DILI severity 0 (no documented liver toxicity). From class II, we chose aspirin (DILI severity 0 but has some potential for liver damage, such as elevated ALT upon prolonged high-dose usage and signs of hepatic injury (Athreya et al., 1973; Barone et al., 1976; Goldenberg, 1974; Koppes and Arnett, 1974; Rich and Johnson, 1973; 1973)), chlorpromazine (DILI severity 2, “adverse reactions” label), clotrimazole (DILI severity 3 and “adverse reactions” label), and ibuprofen (DILI severity 3 and “warnings and precautions” label). From class III, we chose atorvastatin (DILI severity 5 and “warnings and precautions” label), cyclosporin A (DILI severity 7 and “warnings and precautions” label), and amiodarone (DILI severity 8 and “box warning” label). Drugs from class IV (Ambiguous-DILI-concern) were disregarded.

**Table 1.**
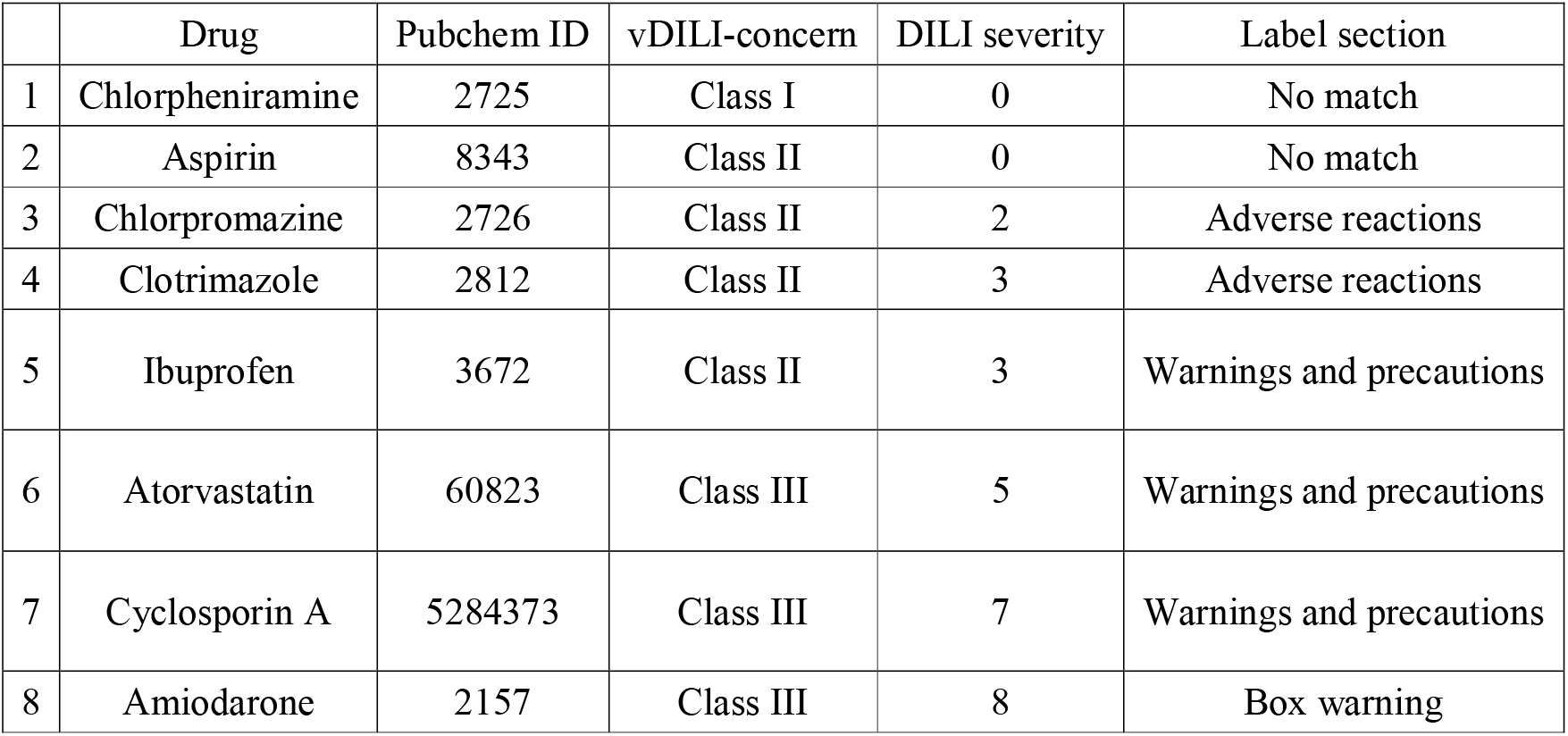
Eight drugs with known hepatotoxicity effects ranked by their v-DILI concern classes, DILI severity scores, and FDA labels.

Next, to obtain information on cell response to the eight drugs *in vitro*, we searched the literature for dose-response studies of the said drugs in human or animal-derived cells (Table 2). We chose studies that had dose-response curves for the derivation of ED50 and ED10 values. In addition, to ensure comparability, we chose studies that evaluated the effects of the drugs specifically after 24-hour exposure (except aspirin and ibuprofen, for which we could not find dose-response studies with 24-hour exposure data). Then, we ranked the drugs by their effect on cell viability according to available data (ED50 and ED10 values). This analysis was carried to compare the effects on human ES cells in the next step.

**Table 2.**
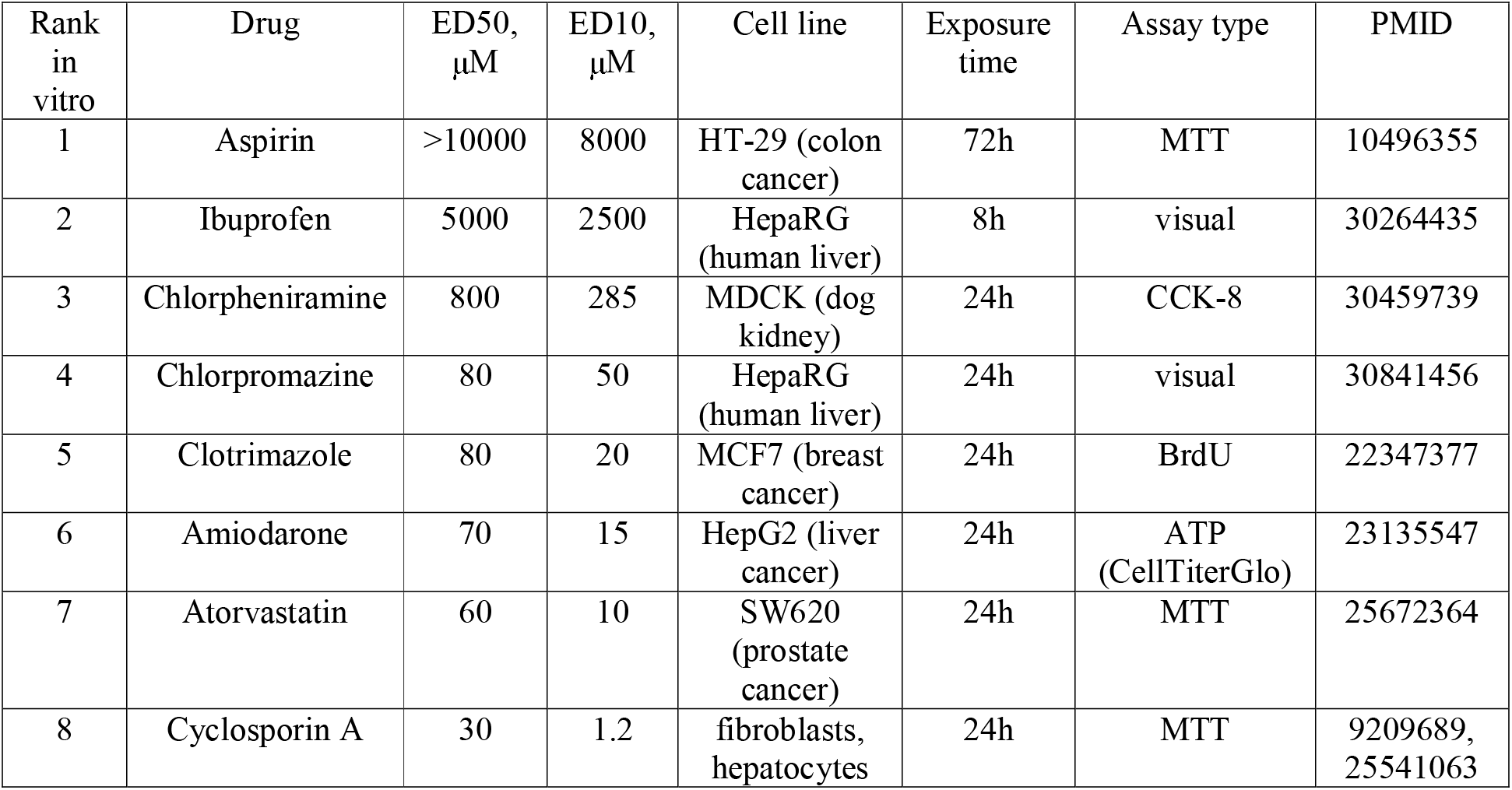
Eight drugs ranked by their effects on cell viability according to literature, where the most effective drug is ranked 8 and the least effective is ranked 1.

### Resistance of human ES cells to selected drugs

Next, we applied the said drugs to human embryonic cell line KhES-3 and obtained the ED10 values as described in Materials and Methods. Figure 1 shows the dose-response curves for the eight drugs, which were obtained by applying the log-logistic function to the ATP cell viability assay data in KhES-3 cell line. Table 3 includes the obtained ED10 values in KhES-3 cell line, the ranks of the drugs in KhES-3 cell line as determined by our assay, the *in vitro* ranks obtained from previous analysis (Table 2), the DILI severity scores, and the vDILI-concern classes. The results reveal a general tendency for the higher ranks of all categories to correspond. Importantly, the correlation coefficient for the *in vitro* results obtained by literature search and for our data in KhES-3 cell line was 0.9, suggesting that the *in vitro* results for ES cells correlate well with those for non-ES cells. On the other hand, there was a marked difference between *in vitro* and *in vivo* categories (*in vitro* ranks obtained in cell lines versus DILI severity scores and vDILI-concern classes, the data of which were obtained from findings in human organism), the correlation coefficient of which was only 0.6, suggesting that *in vitro* results cannot be directly translated to human liver toxicity. Interestingly, for all of the drugs analyzed, the ED10 values were lower in ES cells than differentiated cell lines, such as HepaRG or MCF7, for which we were able to obtain ED10 values from the literature (see Table 2 for cell line names and sources). The biggest difference was observed for chlorpromazine, where ES cells were 16 times more sensitive to the drug than differentiated cells, in this case HepaRG. ES cells showed higher sensitivity to all of the drugs than the other cell types.

**Fig. 1.**
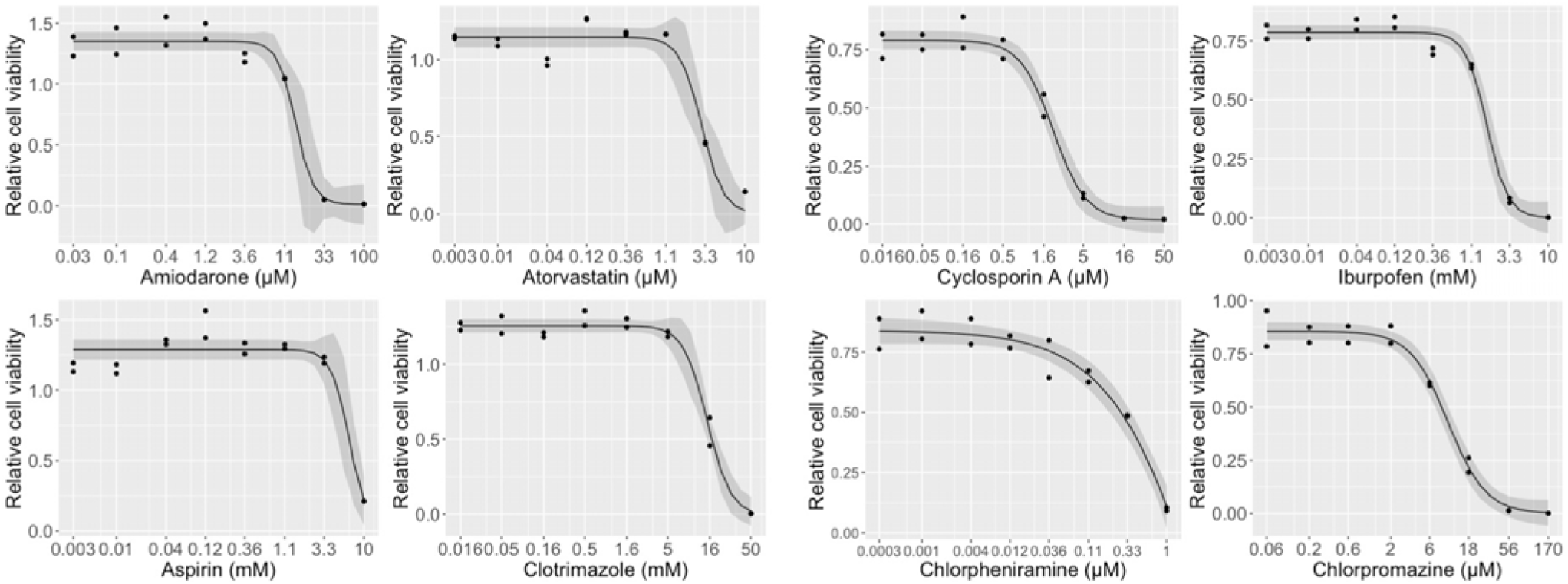
Dose-response curves obtained from ATP assay results using KhES-3 cell line and eight chosen drugs. The black line is obtained by fitting the log-logistic function from the nonlinear regression model provided by *drc* package (Ritz et al., 2015), in which the independent variable is the concentration of the drug and the dependent variable is cell viability. Confidence interval (95%) is depicted in gray.

**Table 3.**
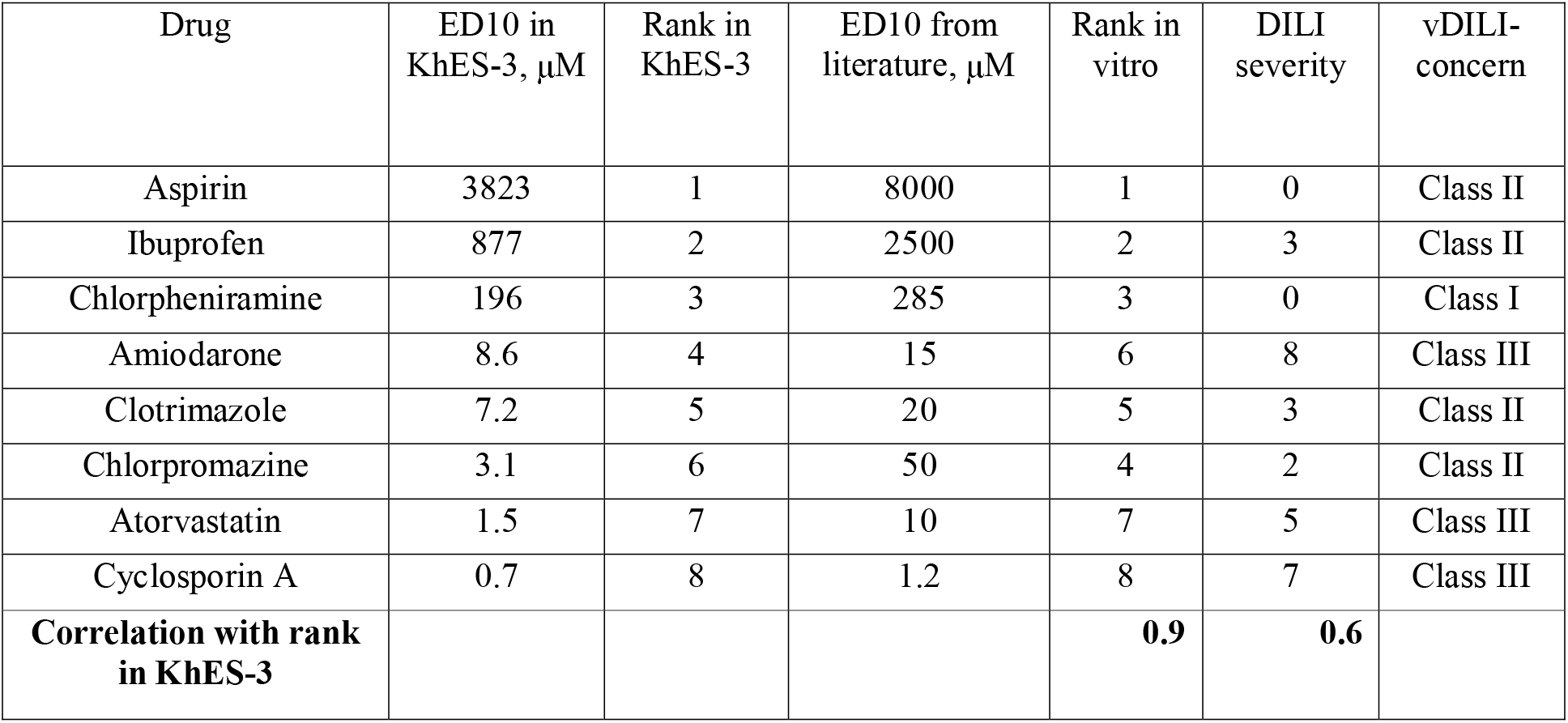
Eight selected drugs and their ED10 values in KhES-3 cell line as determined by the ATP assay, ranks in KhES-3 cell line, ranks in hepatocytes or other cell lines *in vitro*, DILI severity scores, vDILI-concern classes.

| Drug | ED10 in KhES-3, $\mu\text{M}$ | Rank in KhES-3 | ED10 from literature, $\mu\text{M}$ | Rank in vitro | DILI severity | vDILI-concern |
| --- | --- | --- | --- | --- | --- | --- |
| Aspirin | 3823 | 1 | 8000 | 1 | 0 | Class II |
| Ibuprofen | 877 | 2 | 2500 | 2 | 3 | Class II |
| Chlorpheniramine | 196 | 3 | 285 | 3 | 0 | Class I |
| Amiodarone | 8.6 | 4 | 15 | 6 | 8 | Class III |
| Clotrimazole | 7.2 | 5 | 20 | 5 | 3 | Class II |
| Chlorpromazine | 3.1 | 6 | 50 | 4 | 2 | Class II |
| Atorvastatin | 1.5 | 7 | 10 | 7 | 5 | Class III |
| Cyclosporin A | 0.7 | 8 | 1.2 | 8 | 7 | Class III |
| **Correlation with rank in KhES-3** |  |  |  | **0.9** | **0.6** |  |

### Comparison of human ES and iPS cells by their resistance to drug exposure

Our next goal was to compare drug resistance of human iPS cells and human ES cells. For this purpose, we used human iPS cell line RIKEN-2A and compared its response with that of KhES-3 cell line. Figure 2 shows the dose-response curves for RIKEN-2A cell line and Table 4 shows the ED10 values for RIKEN-2A and KhES-3 cell lines. We chose the four drugs in the following fashion: aspirin as the lowest-potency drug, atorvastatin as the highest-potency drug (we did not select cyclosporin A because there are multiple reports that it generates an abnormal dose-response curve due to the stimulation of cell proliferation at low doses), and two drugs in between (amiodarone and clotrimazole). The results demonstrate that the overall difference in ED10 values did not exceed 1.6 times, indicating that the sensitivity to drug exposure did not significantly differ between iPS cells and ES cells.

**Fig. 2.**
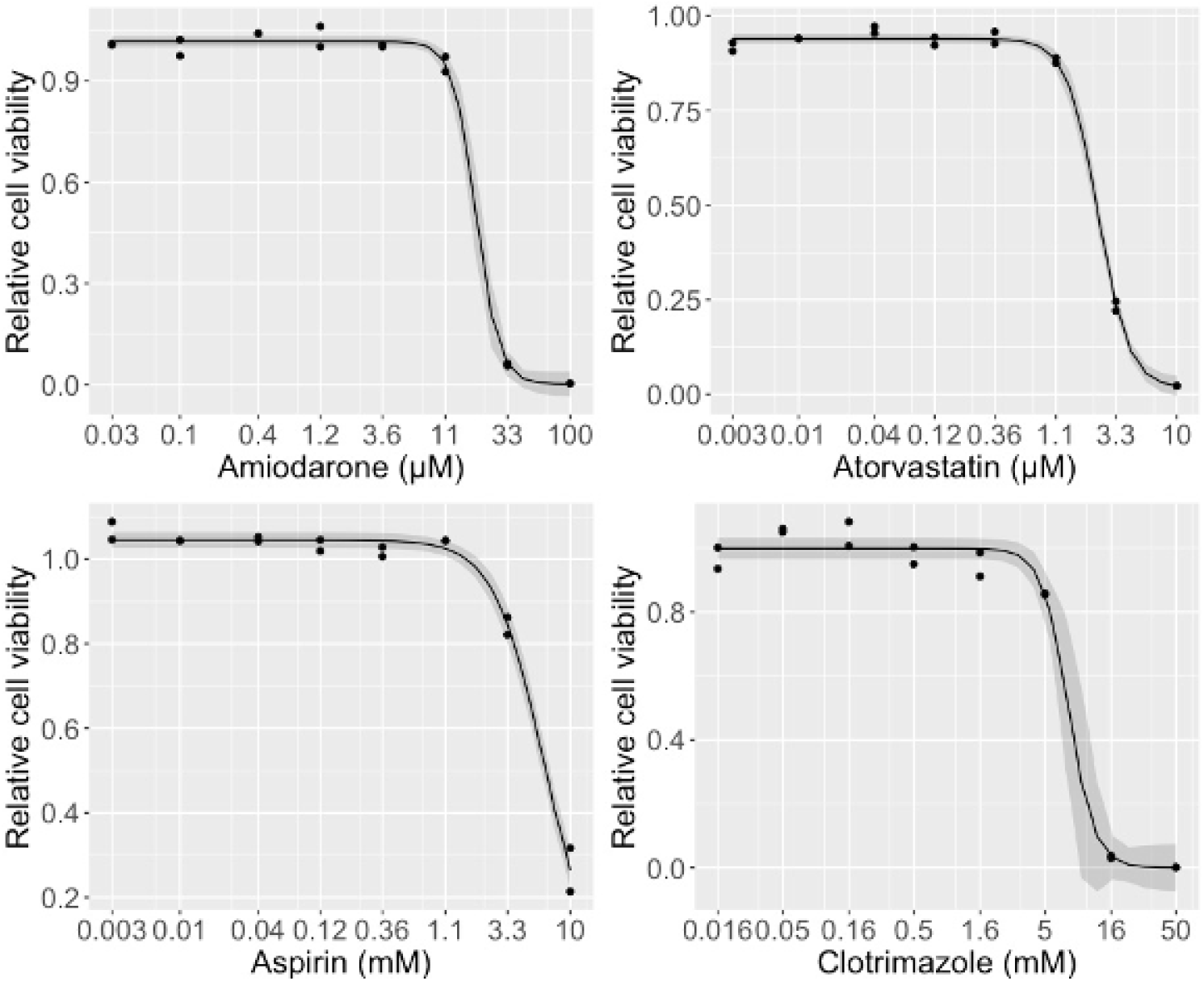
Dose-response curves obtained from ATP assay results using RIKEN-2A cell line and four chosen drugs. The black line is obtained by fitting the log-logistic function from the nonlinear regression model provided by *drc* package (Ritz et al., 2015), in which the independent variable is the concentration of the drug and the dependent variable is cell viability. Confidence interval (95%) is depicted in gray.

**Table 4.**
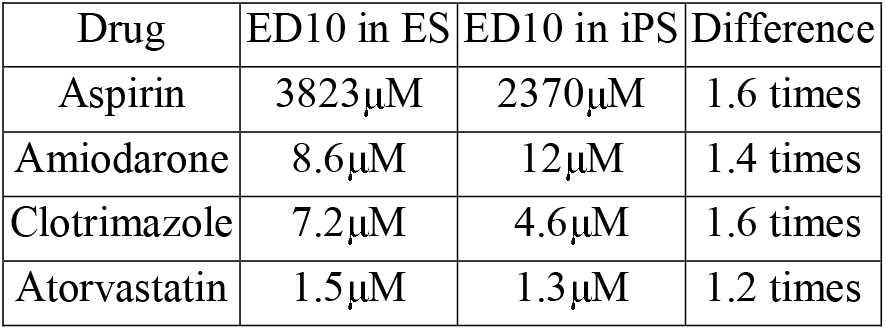
Four selected drugs and their ED10 values in two human cell lines, ES cell line KhES-3 and iPS cell line RIKEN-2A, as determined by the ATP assay.

| Drug | ED10 in ES | ED10 in iPS | Difference |
| --- | --- | --- | --- |
| Aspirin | 3823$\mu\text{M}$ | 2370$\mu\text{M}$ | 1.6 times |
| Amiodarone | 8.6$\mu\text{M}$ | 12$\mu\text{M}$ | 1.4 times |
| Clotrimazole | 7.2$\mu\text{M}$ | 4.6$\mu\text{M}$ | 1.6 times |
| Atorvastatin | 1.5$\mu\text{M}$ | 1.3$\mu\text{M}$ | 1.2 times |

### Establishment of EOS stable cell line with “high-quality” iPS cells

Next, to manipulate the “degree of pluripotency of the cells” (i.e., “quality of iPS cells” (as defined in (Hotta et al., 2009a, 2009b)), we exploited the EOS system. To obtain a more homogeneous and more naïve-like population of cells, we introduced the *piggyBac* based EOS-C(3+)-GFP/puroR vector (PB-EOS), as previously described (Hotta et al., 2009b; Takashima et al., 2014). The expression of this reporter is driven by mouse regulatory elements that are said to be active in undifferentiated ES cells: a trimer of the CR4 element from the Oct3/4 (Pou5f1) distal enhancer and the early transposon (ETn) long terminal repeat promoter. The PB-EOS reporter was shown to be upregulated during naïve-state induction and to maintain visible expression in naïve-like cells (Takashima et al., 2014). We have established stable cell line RIKEN-2A-EOS that is resistant to puromycin, as described in Materials and Methods. Upon introduction of the vector, we found that the resultant line had different populations of GFP-expressing cells, namely, “bright GFP cells” and “dim GFP cells”. Because we observed this phenomenon in the four cell lines we tested, namely, KhES-1 human ES cells, KhES-3 human ES cells, RIKEN-1A human iPS cells, and RIKEN-2A human iPS cells, we think it is an intrinsic property of EOS in pluripotent cell lines (Sup. Fig. 2). As puromycin expression is coupled to GFP expression by the IRES insert in EOS vector, all puromycin-resistant cells, by definition, express GFP with the Oct4 distal enhancer driving the expression. Therefore, GFP expression and puromycin resistance are indicators of the naïve-like state, and all puromycin-resistant cells should possess certain qualities of naïve cells. The heterogeneity of GFP expression and the differences between “bright” and “dim” populations need further investigation to clarify the exact differences between them and are beyond the scope of this study.

### Application of naïve-state induction methods and effect on drug sensitivity

To further advance the cells into the “naïve” state, we tried different methods of naïve-state induction (Sup. Fig. 1). After trying different conditions, we found the highest percentage of round colonies with the highest increase in naïve-state markers upon applying a modified version of the YAP method. Round colonies obtained from RIKEN-2A-EOS-YAP cells stained positive for naïve-state markers SUSD2 and KLF17, and were GFP-positive (Sup. Fig. 3). We next used the ATP assay to investigate the differences in drug resistance between RIKEN-2A-EOS-YAP cells and their progeny obtained after several passages of this cell line in non-YAP medium (AK02N). We performed the ATP assay on day 0 (RIKEN-2A-EOS-YAP immediately after establishment, in YAP medium) and passage days 2, 5, and 10 (RIKEN-2A-EOS-YAP P2, P5, and P10) in AK02N medium. The results are shown in Fig. 3. RIKEN-2A-EOS-YAP cells displayed significantly lower drug resistance than original RIKEN-2A cells and were extremely sensitive to drug exposure both in YAP medium (P0) and after two passages in AK02N medium (P2). Interestingly, after five passages, drug resistance returned and reached pre-naïve levels (Fig. 3 and Table 5). We conclude that YAP treatment of EOS-selected cells produced a sharp decrease in overall drug resistance, which subsided upon returning the cells into AK02N medium and passaging them for several days.

**Fig. 3.**
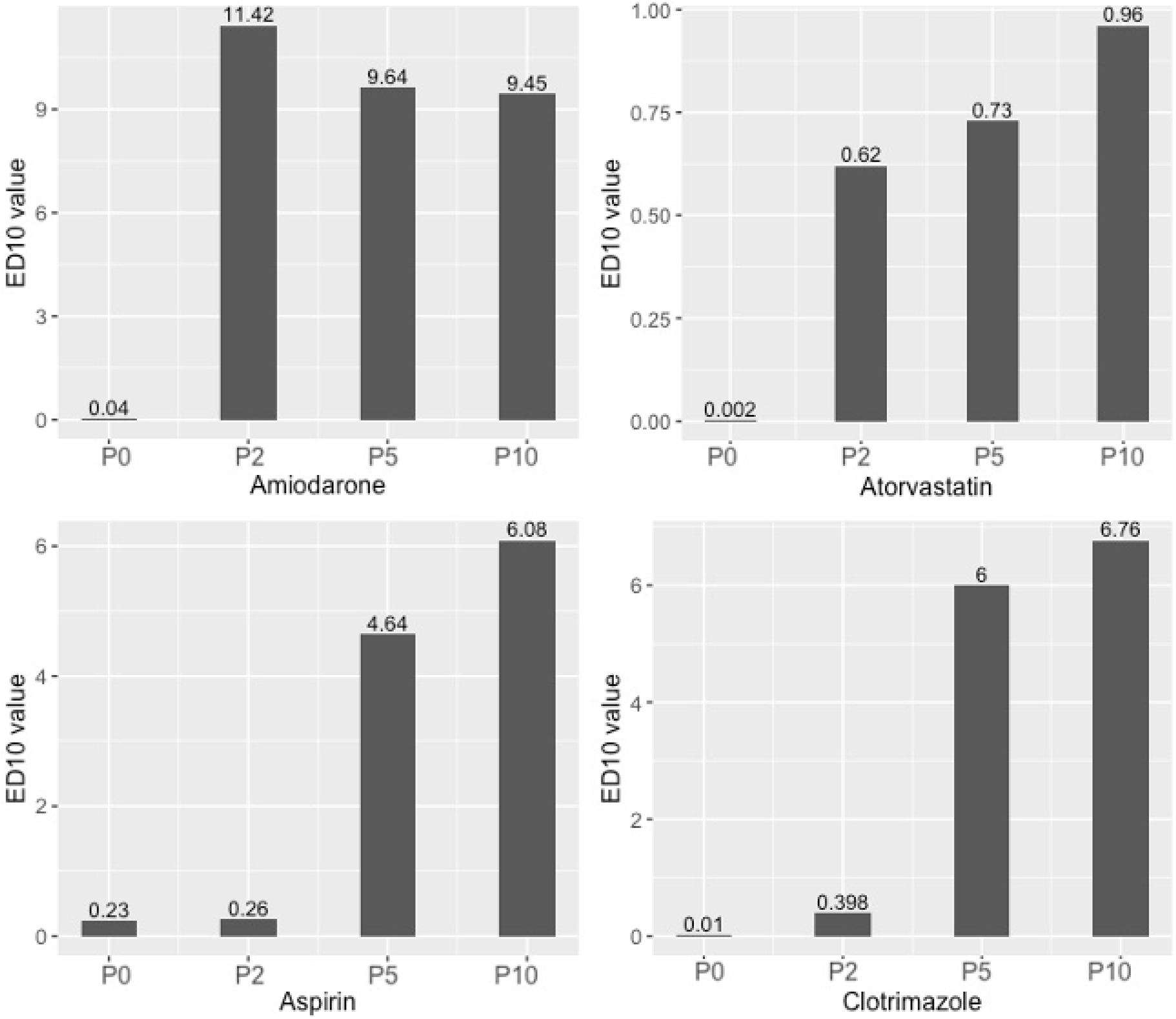
Changes in sensitivity to drug exposure in RIKEN-2A-EOS-YAP cell line with passages in AK02N medium. The following four drugs were used to assess viability changes: aspirin, amiodarone, atorvastatin, and clotrimazole. Passage is indicated on the x-axis, and ED10 value in millimolar for aspirin, and in micromolar for other drugs, is indicated on the y-axis. EOS-YAP cells (Passage 0) display extreme sensitivity to most drugs. Their sensitivity to aspirin, although greatly reduced, is still in the millimolar order, with ED10 = 0.23 *milli*molar, as opposed to micromolar concentrations for the other drugs. At Passage 2, cells regain their relative drug resistance for amiodarone and atorvastatin, but not for aspirin or clotrimazole, for which the ED10 values remain approximately the same as those at Passage 0. At Passage 5, all cells regain their relative drug resistance with ED10 values resembling those of original iPS cell line RIKEN-2A.

**Table 5.** Four selected drugs and their ED10 values in human iPS cell line RIKEN-2A subjected to EOS selection, YAP treatment, and passaging in AK02N medium as determined by the ATP assay.

|  | RIKEN-2A-EOS-YAP | RIKEN-2A-EOS-YAP P2 | RIKEN-2A-EOS-YAP P5 | RIKEN-2A-EOS-YAP P10 |
| --- | --- | --- | --- | --- |
| Amiodarone, $\mu\text{M}$ | 0.04 | 11.42 | 9.64 | 9.45 |
| Atorvastatin, $\mu\text{M}$ | 0.002 | 0.62 | 0.73 | 0.96 |
| Aspirin, mM | 0.23 | 0.26 | 4.64 | 6.08 |
| Clotrimazole, $\mu\text{M}$ | 0.01 | 0.40 | 6.00 | 6.76 |

## Discussion

Human PSCs are an invaluable tool for basic research, clinical applications, and toxicological studies. Nevertheless, the toxicological properties of undifferentiated human PSCs have not been properly investigated to date, and information on the drug resistance of different cell lines in different states of pluripotency is lacking. In this work, we investigated the drug sensitivity of human ES cells, iPS cells, and iPS cells cultured under EOS selection and naïve-state induction conditions. At the first stage of our work, we carefully selected eight drugs having various degrees of toxicity to human organism (specifically, the liver), and included all DILIrank categories of toxicity in our investigation. Then, we exposed human ES cell line KhES-3 to selected drugs and compared the response by rankings and, where possible, by known ED10 values, and found that ES cells possessed lower drug resistance than differentiated cells. Moreover, our *in vitro* results correlated well with *in vitro* results obtained from literature, whereas *in vivo* DILI data did not correlate well with *in vitro* data. It is important to note that currently, there is a lack of sufficient evidence that *in vitro* data from experiments using cell lines could be translated to the *in vivo* effects of drugs on the human body. In fact, our results in Table 3, specifically the correlation between *in vitro* toxicity ranks (Table 3, columns 3 and 5) and *in vivo* DILI toxicity ranks (Table 3, column 6), suggest that there is little correlation between *in vitro* and *in vivo* toxicities. We thus highlight that at present, hepatotoxicity (drug-induced liver injury, DILI) can only be properly assessed by *in vivo* experiments.

Next, we compared human iPS cell line RIKEN-2A cultured under the same conditions as KhES-3, with KhES-3 cells. The ED10 values of the four tested drugs in iPS cells did not significantly differ from the ED10 values in ES cells. We conclude that RIKEN-2A iPS cells possessed approximately the same drug resistance properties as KhES-3 cell line. Then, we exploited the EOS vector for the specific selection of naïve-like cells, established the stable cell line RIKEN-2A-EOS, and confirmed that this cell line shows resistance to puromycin and expresses GFP. In addition, we applied a modified YAP naïve-state induction method to achieve a more homogeneous pluripotent state. The round colonies with clearly defined borders obtained from RIKEN-2A-EOS-YAP cells stained positive for naïve-state markers SUSD2 and KLF17 (Sup. Fig. 3). On comparing the drug resistance of RIKEN-2A-EOS-YAP cells with that of parental RIKEN-2A cells, we found that EOS-YAP modified cells exhibited much lower drug resistance. This sensitivity to drug exposure decreased upon returning the cells into the AK02N medium and passaging them for several days.

It is tempting to speculate that the sharp decrease in drug resistance is due to differences in cell medium as RIKEN-2A-EOS-YAP cells were kept in YAP medium, which is substantially different from the AK02N medium. However, the results in Fig. 3 show a consistent pattern of gradual decrease in drug sensitivity of the cells upon returning them into the AK02N medium. This gradual decrease cannot be explained by medium differences because P2, P5, and P10 cells were kept in the same medium and still retained substantially high drug sensitivity that decreased only by passaging. This suggests that YAP treatment induced a change in intrinsic properties of the cells themselves, such as changes in gene or protein expression.

Another important point to note is that our RIKEN-2A-EOS-YAP cells should not be equated with perfect “naïve iPS” cells as we have not performed the whole variety of experiments needed to confirm the naïve state, such as immunostaining with multiple antibodies (KLF2/4, PECAM1, TFE3 (Betschinger et al., 2013; Takashima et al., 2014)), RNA-seq analysis, etc. Our immunostaining results for SUSD2 and KLF17 can be considered inconclusive. However, it should be noted that the goal of this study was not to produce perfect naïve cells but to investigate the influence of different pluripotent cell states on the drug sensitivity of the cells. To this end, we have achieved this goal and revealed that iPS cells possessed roughly the same drug resistance as ES cells, whereas naïve-state induction cells were significantly different (much less resistant to the drugs). We have also shown that drug resistance returned to pre-naïve levels upon returning the cells into the AK02N medium.

Overall, our study provides important, currently lacking information on drug resistance in iPS and ES cells. This study should serve as an opening to investigations of the drug sensitivity of pluripotent cells and encourage more research on the correlation between stemness and drug resistance.

## Acknowledgements

We thank Professor Akitsu Hotta for kindly providing the PB-EOS vector and explanations.

## References

Adler, S., Pellizzer, C., Hareng, L., Hartung, T., and Bremer, S. (2008). First steps in establishing a developmental toxicity test method based on human embryonic stem cells. Toxicol. In Vitro 22, 200–211.

Ambudkar, S.V., Dey, S., Hrycyna, C.A., Ramachandra, M., Pastan, I., and Gottesman, M.M. (1999). Biochemical, cellular, and pharmacological aspects of the multidrug transporter. Annu. Rev. Pharmacol. Toxicol. 39, 361–398.

Athreya, B.H., Gorske, A.L., and Myers, A.R. (1973). Aspirin-induced abnormalities of liver function. Am J Dis Child 126, 638–641.

Barone, R., Chase, P.H., and Wallace, S.L. (1976). Letter: Salicylate-induced hepatic injury. Arthritis Rheum. 19, 964–966.

Betschinger, J., Nichols, J., Dietmann, S., Corrin, P.D., Paddison, P.J., and Smith, A. (2013). Exit from pluripotency is gated by intracellular redistribution of the bHLH transcription factor Tfe3. Cell 153, 335–347.

Brons, I.G.M., Smithers, L.E., Trotter, M.W.B., Rugg-Gunn, P., Sun, B., Chuva de Sousa Lopes, S.M., Howlett, S.K., Clarkson, A., Ahrlund-Richter, L., Pedersen, R.A., et al. (2007). Derivation of pluripotent epiblast stem cells from mammalian embryos. Nature 448, 191–195.

Carter, M.G., Smagghe, B.J., Stewart, A.K., Rapley, J.A., Lynch, E., Bernier, K.J., Keating, K.W., Hatziioannou, V.M., Hartman, E.J., and Bamdad, C.C. (2016). A primitive growth factor, NME7AB, is sufficient to induce stable naïve state human pluripotency; reprogramming in this novel growth factor confers superior differentiation. Stem Cells 34, 847–859.

Chan, Y.-S., Göke, J., Ng, J.-H., Lu, X., Gonzales, K.A.U., Tan, C.-P., Tng, W.-Q., Hong, Z.-Z., Lim, Y.-S., and Ng, H.-H. (2013). Induction of a human pluripotent state with distinct regulatory circuitry that resembles preimplantation epiblast. Cell Stem Cell 13, 663–675.

Chen, H., Aksoy, I., Gonnot, F., Osteil, P., Aubry, M., Hamela, C., Rognard, C., Hochard, A., Voisin, S., Fontaine, E., et al. (2015). Reinforcement of STAT3 activity reprogrammes human embryonic stem cells to naive-like pluripotency. Nat. Commun. 6, 7095.

Chen, M., Suzuki, A., Thakkar, S., Yu, K., Hu, C., and Tong, W. (2016). DILIrank: the largest reference drug list ranked by the risk for developing drug-induced liver injury in humans. Drug Discov. Today 21, 648–653.

Choi, H.W., Joo, J.Y., Hong, Y.J., Kim, J.S., Song, H., Lee, J.W., Wu, G., Schöler, H.R., and Do, J.T. (2016). Distinct enhancer activity of oct4 in naive and primed mouse pluripotency. Stem Cell Rep. 7, 911–926.

Christodoulou, C., and Kotton, D.N. (2012). Are embryonic stem and induced pluripotent stem cells the same or different? Implications for their potential therapeutic use. Cell Cycle 11, 5–6.

Duggal, G., Warrier, S., Ghimire, S., Broekaert, D., Van der Jeught, M., Lierman, S., Deroo, T., Peelman, L., Van Soom, A., Cornelissen, R., et al. (2015). Alternative routes to induce naïve pluripotency in human embryonic stem cells. Stem Cells 33, 2686–2698.

Fukunaga-Kalabis, M., and Herlyn, M. (2012). Beyond ABC: another mechanism of drug resistance in melanoma side population. J. Invest. Dermatol. 132, 2317–2319.

Gillet, J.-P., and Gottesman, M.M. (2010). Mechanisms of multidrug resistance in cancer. Methods Mol. Biol. 596, 47–76.

Goldenberg, D.L. (1974). Letter: Aspirin hepatotoxicity. Ann. Intern. Med. 80, 773.

Gottesman, M.M., Fojo, T., and Bates, S.E. (2002). Multidrug resistance in cancer: role of ATP-dependent transporters. Nat. Rev. Cancer 2, 48–58.

Greber, B., Wu, G., Bernemann, C., Joo, J.Y., Han, D.W., Ko, K., Tapia, N., Sabour, D., Sterneckert, J., Tesar, P., et al. (2010). Conserved and divergent roles of FGF signaling in mouse epiblast stem cells and human embryonic stem cells. Cell Stem Cell 6, 215–226.

Gu, Q., Hao, J., Zhao, X., Li, W., Liu, L., Wang, L., Liu, Z., and Zhou, Q. (2012). Rapid conversion of human ESCs into mouse ESC-like pluripotent state by optimizing culture conditions. Protein Cell 3, 71–79.

Hackett, C.H., and Fortier, L.A. (2011). Embryonic stem cells and iPS cells: sources and characteristics. Vet Clin North Am Equine Pract 27, 233–242.

Hanna, J., Cheng, A.W., Saha, K., Kim, J., Lengner, C.J., Soldner, F., Cassady, J.P., Muffat, J., Carey, B.W., and Jaenisch, R. (2010). Human embryonic stem cells with biological and epigenetic characteristics similar to those of mouse ESCs. Proc. Natl. Acad. Sci. USA 107, 9222–9227.

Hotta, A., Cheung, A.Y.L., Farra, N., Garcha, K., Chang, W.Y., Pasceri, P., Stanford, W.L., and Ellis, J. (2009a). EOS lentiviral vector selection system for human induced pluripotent stem cells. Nat. Protoc. 4, 1828–1844.

Hotta, A., Cheung, A.Y.L., Farra, N., Vijayaragavan, K., Séguin, C.A., Draper, J.S., Pasceri, P., Maksakova, I.A., Mager, D.L., Rossant, J., et al. (2009b). Isolation of human iPS cells using EOS lentiviral vectors to select for pluripotency. Nat. Methods 6, 370–376.

Hu, L., McArthur, C., and Jaffe, R.B. (2010). Ovarian cancer stem-like side-population cells are tumourigenic and chemoresistant. Br. J. Cancer 102, 1276–1283.

Keller, G. (2005). Embryonic stem cell differentiation: emergence of a new era in biology and medicine. Genes Dev. 19, 1129–1155.

Koppes, G.M., and Arnett, F.C. (1974). Salicylate hepatotoxicity. Postgrad. Med. 56, 193–195.

Kubara, K., Yamazaki, K., Ishihara, Y., Naruto, T., Lin, H.-T., Nishimura, K., Ohtaka, M., Nakanishi, M., Ito, M., Tsukahara, K., et al. (2018). Status of KRAS in iPSCs Impacts upon Self-Renewal and Differentiation Propensity. Stem Cell Rep. 11, 380–394.

Kusakawa, S., Yamauchi, J., Miyamoto, Y., Sanbe, A., and Tanoue, A. (2008). Estimation of embryotoxic effect of fluoxetine using embryonic stem cell differentiation system. Life Sci. 83, 871–877.

Laschinski, G., Vogel, R., and Spielmann, H. (1991). Cytotoxicity test using blastocyst-derived euploid embryonal stem cells: a new approach to in vitro teratogenesis screening. Reprod Toxicol 5, 57–64.

Li, D., Su, D., Xue, L., Liu, Y., and Pang, W. (2015). Establishment of pancreatic cancer stem cells by flow cytometry and their biological characteristics. Int J Clin Exp Pathol 8, 11218–11223.

Li, W., Wei, W., Zhu, S., Zhu, J., Shi, Y., Lin, T., Hao, E., Hayek, A., Deng, H., and Ding, S. (2009). Generation of rat and human induced pluripotent stem cells by combining genetic reprogramming and chemical inhibitors. Cell Stem Cell 4, 16–19.

Liu, H., Ren, C., Liu, W., Jiang, X., Wang, L., Zhu, B., Jia, W., Lin, J., Tan, J., and Liu, X. (2017a). Embryotoxicity estimation of commonly used compounds with embryonic stem cell test. Mol. Med. Rep. 16, 263–271.

Liu, X., Nefzger, C.M., Rossello, F.J., Chen, J., Knaupp, A.S., Firas, J., Ford, E., Pflueger, J., Paynter, J.M., Chy, H.S., et al. (2017b). Comprehensive characterization of distinct states of human naive pluripotency generated by reprogramming. Nat. Methods 14, 1055–1062.

Luo, Y., Ellis, L.Z., Dallaglio, K., Takeda, M., Robinson, W.A., Robinson, S.E., Liu, W., Lewis, K.D., McCarter, M.D., Gonzalez, R., et al. (2012). Side population cells from human melanoma tumors reveal diverse mechanisms for chemoresistance. J. Invest. Dermatol. 132, 2440–2450.

MacDonagh, L., Gray, S.G., Breen, E., Cuffe, S., Finn, S.P., O’Byrne, K.J., and Barr, M.P. (2016). Lung cancer stem cells: The root of resistance. Cancer Lett. 372, 147–156.

Martin, G.R. (1981). Isolation of a pluripotent cell line from early mouse embryos cultured in medium conditioned by teratocarcinoma stem cells. Proc. Natl. Acad. Sci. USA 78, 7634–7638.

Narsinh, K.H., Plews, J., and Wu, J.C. (2011). Comparison of human induced pluripotent and embryonic stem cells: fraternal or identical twins? Mol. Ther. 19, 635–638.

Niess, H., Camaj, P., Renner, A., Ischenko, I., Zhao, Y., Krebs, S., Mysliwietz, J., Jäckel, C., Nelson, P.J., Blum, H., et al. (2015). Side population cells of pancreatic cancer show characteristics of cancer stem cells responsible for resistance and metastasis. Target Oncol 10, 215–227.

Ochiai, H., Suga, H., Yamada, T., Sakakibara, M., Kasai, T., Ozone, C., Ogawa, K., Goto, M., Banno, R., Tsunekawa, S., et al. (2015). BMP4 and FGF strongly induce differentiation of mouse ES cells into oral ectoderm. Stem Cell Res. 15, 290–298.

Onishi, K., and Zandstra, P.W. (2015). LIF signaling in stem cells and development. Development 142, 2230–2236.

Panova, A.V., Bogomazova, A.N., Lagarkova, M.A., and Kiselev, S.L. (2018). Epigenetic reprogramming by naïve conditions establishes an irreversible state of partial X chromosome reactivation in female stem cells. Oncotarget 9, 25136–25147.

Park, T.S., Zimmerlin, L., Evans-Moses, R., and Zambidis, E.T. (2018). Chemical Reversion of Conventional Human Pluripotent Stem Cells to a Naïve-like State with Improved Multilineage Differentiation Potency. J. Vis. Exp.

Qin, H., Hejna, M., Liu, Y., Percharde, M., Wossidlo, M., Blouin, L., Durruthy-Durruthy, J., Wong, P., Qi, Z., Yu, J., et al. (2016). YAP induces human naive pluripotency. Cell Rep. 14, 2301–2312.

Rich, R.R., and Johnson, J.S. (1973). Salicylate hepatotoxicity in patients with juvenile rheumatoid arthritis. Arthritis Rheum. 16, 1–9.

Ritz, C., Baty, F., Streibig, J.C., and Gerhard, D. (2015). Dose-Response Analysis Using R. PLoS One 10, e0146021.

Shen, D.W., Goldenberg, S., Pastan, I., and Gottesman, M.M. (2000). Decreased accumulation of [14C]carboplatin in human cisplatin-resistant cells results from reduced energy-dependent uptake. J. Cell Physiol. 183, 108–116.

Takahashi, K., and Yamanaka, S. (2006). Induction of pluripotent stem cells from mouse embryonic and adult fibroblast cultures by defined factors. Cell 126, 663–676.

Takashima, Y., Guo, G., Loos, R., Nichols, J., Ficz, G., Krueger, F., Oxley, D., Santos, F., Clarke, J., Mansfield, W., et al. (2014). Resetting transcription factor control circuitry toward ground-state pluripotency in human. Cell 158, 1254–1269.

Tandon, S., and Jyoti, S. (2012). Embryonic stem cells: An alternative approach to developmental toxicity testing. J. Pharm. Bioallied Sci. 4, 96–100.

Tesar, P.J., Chenoweth, J.G., Brook, F.A., Davies, T.J., Evans, E.P., Mack, D.L., Gardner, R.L., and McKay, R.D.G. (2007). New cell lines from mouse epiblast share defining features with human embryonic stem cells. Nature 448, 196–199.

Theunissen, T.W., Powell, B.E., Wang, H., Mitalipova, M., Faddah, D.A., Reddy, J., Fan, Z.P., Maetzel, D., Ganz, K., Shi, L., et al. (2014). Systematic identification of culture conditions for induction and maintenance of naive human pluripotency. Cell Stem Cell 15, 471–487.

Tsunekuni, K., Konno, M., Haraguchi, N., Koseki, J., Asai, A., Matsuoka, K., Kobunai, T., Takechi, T., Doki, Y., Mori, M., et al. (2019). CD44/CD133-Positive Colorectal Cancer Stem Cells are Sensitive to Trifluridine Exposure. Sci. Rep. 9, 14861.

Valamehr, B., Robinson, M., Abujarour, R., Rezner, B., Vranceanu, F., Le, T., Medcalf, A., Lee, T.T., Fitch, M., Robbins, D., et al. (2014). Platform for induction and maintenance of transgene-free hiPSCs resembling ground state pluripotent stem cells. Stem Cell Rep. 2, 366–381.

Wang, W., Yang, J., Liu, H., Lu, D., Chen, X., Zenonos, Z., Campos, L.S., Rad, R., Guo, G., Zhang, S., et al. (2011). Rapid and efficient reprogramming of somatic cells to induced pluripotent stem cells by retinoic acid receptor gamma and liver receptor homolog 1. Proc. Natl. Acad. Sci. USA 108, 18283–18288.

Ware, C.B., Nelson, A.M., Mecham, B., Hesson, J., Zhou, W., Jonlin, E.C., Jimenez-Caliani, A.J., Deng, X., Cavanaugh, C., Cook, S., et al. (2014). Derivation of naive human embryonic stem cells. Proc. Natl. Acad. Sci. USA 111, 4484–4489.

Warrier, S., Popovic, M., Van der Jeught, M., and Heindryckx, B. (2016). Establishment and characterization of naïve pluripotency in human embryonic stem cells. Methods Mol. Biol. 1516, 13–46.

Warrier, S., Van der Jeught, M., Duggal, G., Tilleman, L., Sutherland, E., Taelman, J., Popovic, M., Lierman, S., Chuva De Sousa Lopes, S., Van Soom, A., et al. (2017). Direct comparison of distinct naive pluripotent states in human embryonic stem cells. Nat. Commun. 8, 15055.

Wouters, J., Stas, M., Gremeaux, L., Govaere, O., Van den Broeck, A., Maes, H., Agostinis, P., Roskams, T., van den Oord, J.J., and Vankelecom, H. (2013). The human melanoma side population displays molecular and functional characteristics of enriched chemoresistance and tumorigenesis. PLoS One 8, e76550.

Yamane, J., Aburatani, S., Imanishi, S., Akanuma, H., Nagano, R., Kato, T., Sone, H., Ohsako, S., and Fujibuchi, W. (2016). Prediction of developmental chemical toxicity based on gene networks of human embryonic stem cells. Nucleic Acids Res. 44, 5515–5528.

Yeh, C.-T., Wu, A.T.H., Chang, P.M.-H., Chen, K.-Y., Yang, C.-N., Yang, S.-C., Ho, C.-C., Chen, C.-C., Kuo, Y.-L., Lee, P.-Y., et al. (2012). Trifluoperazine, an antipsychotic agent, inhibits cancer stem cell growth and overcomes drug resistance of lung cancer. Am. J. Respir. Crit. Care Med. 186, 1180–1188.

Yeom, Y.I., Fuhrmann, G., Ovitt, C.E., Brehm, A., Ohbo, K., Gross, M., Hübner, K., and Schöler, H.R. (1996). Germline regulatory element of Oct-4 specific for the totipotent cycle of embryonal cells. Development 122, 881–894.

Zhou, Y., Xia, L., Wang, H., Oyang, L., Su, M., Liu, Q., Lin, J., Tan, S., Tian, Y., Liao, Q., et al. (2018). Cancer stem cells in progression of colorectal cancer. Oncotarget 9, 33403–33415.

Zimmerlin, L., Park, T.S., Huo, J.S., Verma, K., Pather, S.R., Talbot, C.C., Agarwal, J., Steppan, D., Zhang, Y.W., Considine, M., et al. (2016). Tankyrase inhibition promotes a stable human naïve pluripotent state with improved functionality. Development 143, 4368–4380.

Zimmerlin, L., Park, T.S., Huo, J.S., Verma, K., Pather, S.R., Talbot, C.C., Agarwal, J., Steppan, D., Zhang, Y.W., Considine, M., et al. (1973). Liver injury by salicylates. Br. Med. J. 2, 732.

