## Supplemental Figures and Text for "Human ES and iPS Cells Display Less Drug Resistance Than Differentiated Cells, and Naïve-State Induction Further Decreases Drug Resistance"

**Wataru Fujibuchi**

Address: Center for iPS Cell Research and Application (CiRA), Kyoto University, 53 Kawahara-cho, Shogoin, Sakyo-ku, Kyoto 606-8507, Japan

***Corresponding author:**

**藤渕 航 Ph.D.**

**所属：**京都大学 iPS 細胞研究所

**住所:** 〒606-8507 京都市左京区聖護院川原町 53

### Supplementary Fig. 1

| 5i/LAF (Theunissen 2014) | t2iLGoY (Takashima 2014) | XAV939 (Zambidis 2016) | YAP signal (Qin 2016) |
|---|---|---|---|
| 1 | 2 | 3 | 4 |
| DMEM/F12:N2 B27 +<br>0.5% KSR<br>1% NEAA, 1mM GlutaMax<br>0.2 mM B-mercaptoethanol | DMEM/F12:N2 B27 +<br>0.5% KSR<br>1% NEAA, 1mM GlutaMax<br>0.2 mM B-mercaptoethanol | DMEM/F12<br>+<br>20% KSR<br>1% NEAA, 1mM GlutaMax<br>0.2 mM B-mercaptoethanol | DMEM/F12:N2 B27 +<br>1% NEAA, 1mM GlutaMax<br>0.2 mM B-mercaptoethanol |
| 10 ng/mL FGF | 10 ng/mL FGF | 10 ng/mL FGF | |
| 50 µg/mL BSA | 50 µg/mL BSA | | |
| 10 µM Y-27632 | 10 µM Y-27632 | 10 µM Y-27632 | 10 µM Y-27632 |
| 10 ng/mL LIF | 10 ng/mL LIF | 10 ng/mL LIF | 10 ng/mL LIF |
| 0.5-1 µM PD0325901 | 0.5-1 µM PD0325901 | 0.5-1 µM PD0325901 | 0.5-1 µM PD0325901 |
| | 1 µM CHIR 99021 | 3 µM CHIR 99021 | 3 µM CHIR 99021 |
| 20 ng/mL Activin A | 50 µg/mL Ascorbic acid | 10 µM Forskolin | 10 µM Forskolin |
| 0.5 µM IM-12 | | | |
| 0.5 µM SB590885 | 2.5 µM Go6983 | 4 µM XAV939 | 10 µM LPA |
| 1 µM WH-4-023 | | 2 µM Purmorphamine | |

| 5i/LAF (Theunissen 2014) | t2iLGoY (Takashima 2014) | XAV939 (Zambidis 2016) | YAP signal (Qin 2016) |
|---|---|---|---|
| 5 | 6 | 7 | 8 |
| TeSR – E8 | TeSR – E8 | TeSR – E8 | TeSR – E8 |
| 10 ng/mL FGF | 10 ng/mL FGF | 10 ng/mL FGF | |
| 50 µg/mL BSA | 50 µg/mL BSA | | |
| 10 µM Y-27632 | 10 µM Y-27632 | 10 µM Y-27632 | 10 µM Y-27632 |
| 10 ng/mL LIF | 10 ng/mL LIF | 10 ng/mL LIF | 10 ng/mL LIF |
| 0.5-1 µM PD0325901 | 0.5-1 µM PD0325901 | 0.5-1 µM PD0325901 | 0.5-1 µM PD0325901 |
| | 1 µM CHIR 99021 | 3 µM CHIR 99021 | 3 µM CHIR 99021 |
| 20 ng/mL Activin A | 50 µg/mL Ascorbic acid | 10 µM Forskolin | 10 µM Forskolin |
| 0.5 µM IM-12 | | | |
| 0.5 µM SB590885 | 2.5 µM Go6983 | 4 µM XAV939 | 10 µM LPA |
| 1 µM WH-4-023 | | 2 µM Purmorphamine | |

**Supplementary Fig. 2.**

KhES1-EOS KhES3-EOS

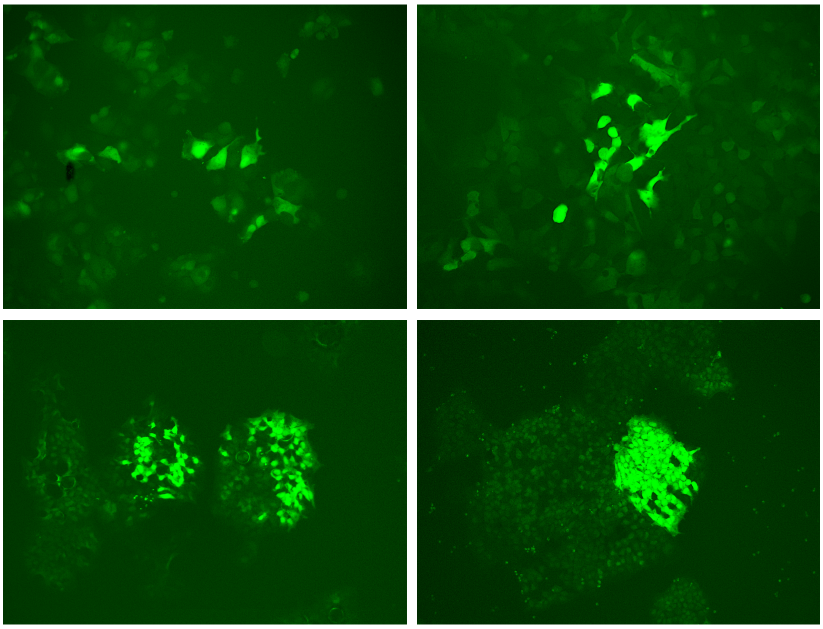

RIKEN1A-EOS RIKEN2A-EOS

**Supplementary Fig. 3.**

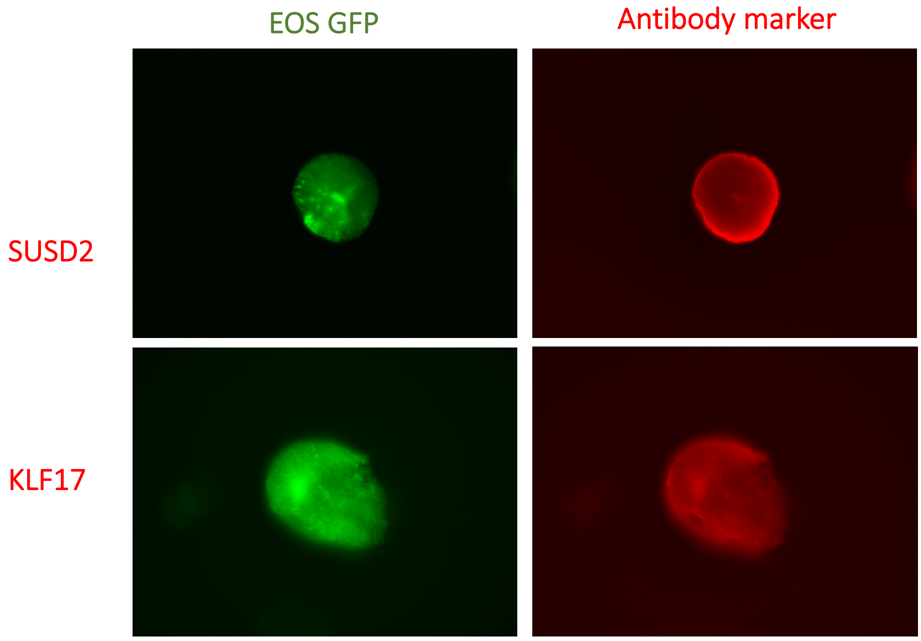

**Supplementary Figure legends**

**Supplementary Fig. 1. Four selected methods of naïve-state induction.**
Four methods (Theunissen et al., 2014, Takashima et al., 2014, Qin et al., 2016, Zimmerlin et al., 2016) were chosen to test naïve-state induction (light blue, upper row). As all four methods can be used with conventional DMEM medium (numbers 1 to 4 in dark blue) and with TeSR-E8 (numbers 5 to 8 in dark blue), eight different conditions were obtained. Medium bases for each corresponding condition are shown in pink, and drug additives, such as FGF or BSA, are shown in green.

**Supplementary Fig. 2. Four cell lines with EOS vector expressing different levels of GFP.**
Four cell lines, as indicated by the names in the figure, were subjected to introduction of (PB) EOS-C(3+)-GFP/puroR vector (EOS), as previously described (Hotta et al., 2009). After puromycin selection, stable cell lines resistant to puromycin were obtained. The cells of the lines expressed GFP in different quantities, roughly dividing the cells of each line into two populations: “bright cells” and “dim cells”. This phenomenon was observed in all four cell lines.

**Supplementary Fig. 3. Immunofluorescence analysis of EOS line RIKEN-2A-EOS subjected to modified YAP naïve-state induction for the presence of naïve-state markers (SUSD2, KLF17).**
Analysis of RIKEN-2A-EOS-YAP line. EOS fluorescence is shown in green and the corresponding markers are shown in red.
